# The genomic health of ancient hominins

**DOI:** 10.1101/145193

**Authors:** Ali J. Berens, Taylor L. Cooper, Joseph Lachance

## Abstract

The genomes of ancient humans, Neandertals, and Denisovans contain many alleles that influence disease risks. Using genotypes at 3180 disease-associated loci, we estimated the disease burden of 147 ancient genomes. After correcting for missing data, genetic risk scores were generated for nine disease categories and the set of all combined diseases. These genetic risk scores were used to examine the effects of different types of subsistence, geography, and sample age on the number of risk alleles in each ancient genome. On a broad scale, hereditary disease risks are similar for ancient hominins and modern-day humans, and the GRS percentiles of ancient individuals span the full range of what is observed in present day individuals. In addition, there is evidence that ancient pastoralists may have had healthier genomes than hunter-gatherers and agriculturalists. We also observed a temporal trend whereby genomes from the recent past are more likely to be healthier than genomes from the deep past. This calls into question the idea that modern lifestyles have caused genetic load to increase over time. Focusing on individual genomes, we find that the overall genomic health of the Altai Neandertal is worse than 97% of present day humans and that Ötzi the Tyrolean Iceman had a genetic predisposition to gastrointestinal and cardiovascular diseases. As demonstrated by this work, ancient genomes afford us new opportunities to diagnose past human health, which has previously been limited by the quality and completeness of remains.

## Introduction

Ancient human remains yield valuable information about the health and disease of individuals from the past. Skeletal remains provide details about medical practices (Andrushko and Verano 2008) and certain diseases, such as cancers (Binder, et al. 2014; Odes, et al. 2016) and rheumatic diseases (Entezami, et al. 2011) that visibly alter or affect bone. However, many diseases do not leave their mark on bone; instead, they cause soft tissue damage requiring mummified remains for diagnosis. One of the best studied set of mummified human remains is the Tyrolean Iceman, a well-preserved 5300-year-old Neolithic man discovered in the Ötztal Alps. Examination of his remains shows hardening arteries suggesting that this man possessed a predisposition for coronary heart disease (Murphy, et al. 2003). Further evidence from the sequencing of the Iceman’s genome supports this diagnosis (Keller, et al. 2012) and demonstrates that modern risk alleles can accurately predict disease risk in ancient individuals. In addition to heart disease, genetic data also indicate that the Iceman was likely lactose intolerant, had type O blood, and had brown eyes (Keller, et al. 2012). Unfortunately, not all ancient human samples are as complete and well-preserved as the Tyrolean Iceman. Genomic information may shed new light on the health of ancient individuals, especially when soft tissue and skeletal evidence are unavailable.

To estimate the health of different individuals, genetic information can be converted into a quantitative measure of hereditary disease burden, which we refer to as a genetic risk score (GRS). GRS have already been used to estimate disease risk in modern humans (Chatterjee, et al. 2016). Assuming that disease loci act independently, one way to calculate GRS is to sum the number of risk alleles across all disease loci after weighting each allele by its effect size (Corona, et al. 2013). The majority of known disease alleles have been identified in present-day Europeans through genome wide association studies (GWAS) using microarrays, which are known to contain a biased set of genetic variants (Lachance and Tishkoff 2013). Despite this bias, genetic risk scores can still be informative about disease risk. For example, GRS for breast, colorectal, and bladder cancer and coronary heart disease accurately predict risk of each disease while offering insights into prevention and treatment options (Chatterjee, et al. 2016). Not only can GRS be applied to specific traits, but genetic risk can also be estimated for the entire genome.

One significant limitation of ancient DNA studies is that most genomes are not complete, which makes it more challenging to estimate disease risk. Genomes may be incomplete due to contamination (Orlando, et al. 2015), DNA degradation (Hofreiter, et al. 2015), or low coverage sequencing, as has been the case with large cohort studies (Mathieson, et al. 2015; Lazaridis, et al. 2016). One approach to address missing genomic data is to impute genotypes at disease-associated loci (Sainz, et al. 1989; Li, Willer, et al. 2009). However, imputation of ancient genotypes is challenging because of small sample sizes. To avoid imputation errors, genetic risk scores for each ancient sample can be calculated using only loci with successful genotype calls (i.e. genetic risk scores for each sample use a different set of disease loci). Each ancient genome can then be compared to modern genomes at a matched set of disease loci to generate standardized GRS percentiles, which enables comparisons between different ancient genomes.

There has been recent debate about whether rates of sequence change in human genomes have increased over time. Hawks, et al. (2007) suggest that rates of adaptive evolution have been accelerating over the last 40,000 years, especially within the last 5,000 years. The holocene has brought new selective pressures associated with the transition from hunting and gathering to agricultural practices (Hawks, et al. 2007). Genes involved in subsistence and dietary changes along with disease resistance genes show strong positive selection (Hawks, et al. 2007; Field, et al. 2016). Others have argued that there has not been enough time for selection to expel deleterious alleles that lead to evolutionary mismatches between our genomes and modern environments (Cordain, et al. 2005). More recently, modern medicine and changes in living conditions may have led to the relaxation of selective pressures within the last century. Along these lines, Lynch (2016) argues that there has been a substantial increase in the mutational load of modern humans (on the order of 1% reduction in fitness per generation). However, Roth and Wakeley (2016) contend that the relaxation of selective pressures is not as severe as suggested by Lynch (2016).

With this debate in mind, our study estimated the genomic health of ancient individuals by combining disease-association information from the NHGRI-EBI GWAS Catalog with publicly available genomes of ancient hominin samples. We investigated relationships between genetic disease risk and sample age. We further explored this dataset by determining whether genetic risks vary for individuals who practiced different types of subsistence and examined whether ancient GRS follow a geographic cline. Finally, we probed differences in genomic health between modern and ancient hominins by calculating genetic risk scores for different disease categories.

## Materials and Methods

### Disease-associated loci

The set of all known autosomal disease-associated alleles was downloaded from the NHGRI-EBI Catalog of published GWAS studies (www.ebi.ac.uk/gwas; last accessed on September 28, 2016) (Burdett, et al. 2016; MacArthur, et al. 2017). Any non-disease-associated alleles (e.g., hair color, blood pressure, and amount of sleep) were removed from consideration. We also filtered out disease-associated alleles from the GWAS Catalog that were missing p-value, odds ratio, and/or allele frequency information. When SNPs were associated with the same disease in multiple GWAS, we retained the SNP with the strongest disease association (i.e. lowest p-value). To ensure independence between variants associated with a given disease, we included only loci that were not in linkage disequilibrium with other disease variants. This involved keeping the strongest disease association (i.e. lowest p-value) whenever two SNPs were located within 100kb of each other. After filtering, we were left with 3180 autosomal disease-associated alleles from 578 GWAS. We classified all remaining disease-associated loci into nine categories (allergy/autoimmune, cancers, cardiovascular, dental/periodontal, gastrointestinal/liver, metabolism/weight, miscellaneous, morphological/muscular, and neurological/psychological). These disease categories are not mutually exclusive, so a single locus could be grouped into multiple categories (e.g., Crohn’s disease is both an autoimmune and a gastrointestinal disease). Note that many disease-associated loci tend are older SNPs that have intermediate frequency alleles (Lachance 2010). Because of this, many GWAS alleles can also be found in Neanderthal and Denisovan genomes.

### Ancient hominin genomes

We downloaded all known publicly available ancient hominin genomes (449 genomes as of July 1, 2016) and identified genotypes at the 3180 focal disease-associated loci. For genomes that were only available as alignments to the human reference genome, we called genotypes from these alignments using the ‘mpileup’ function in SAMtools (v. 1.3) (Li, Handsaker, et al. 2009; Li 2011). We retained ancient hominin samples that had genotype calls at greater than fifty percent of the focal 3180 disease-associated loci, after which 147 ancient genomes from 22 studies remained. This filter was used to ensure sufficient genotype information for subsequent analyses. Figure 1 displays the geographic locations of the 147 focal ancient individuals, which were jittered to clarify densely sampled locations.

**Figure 1:**
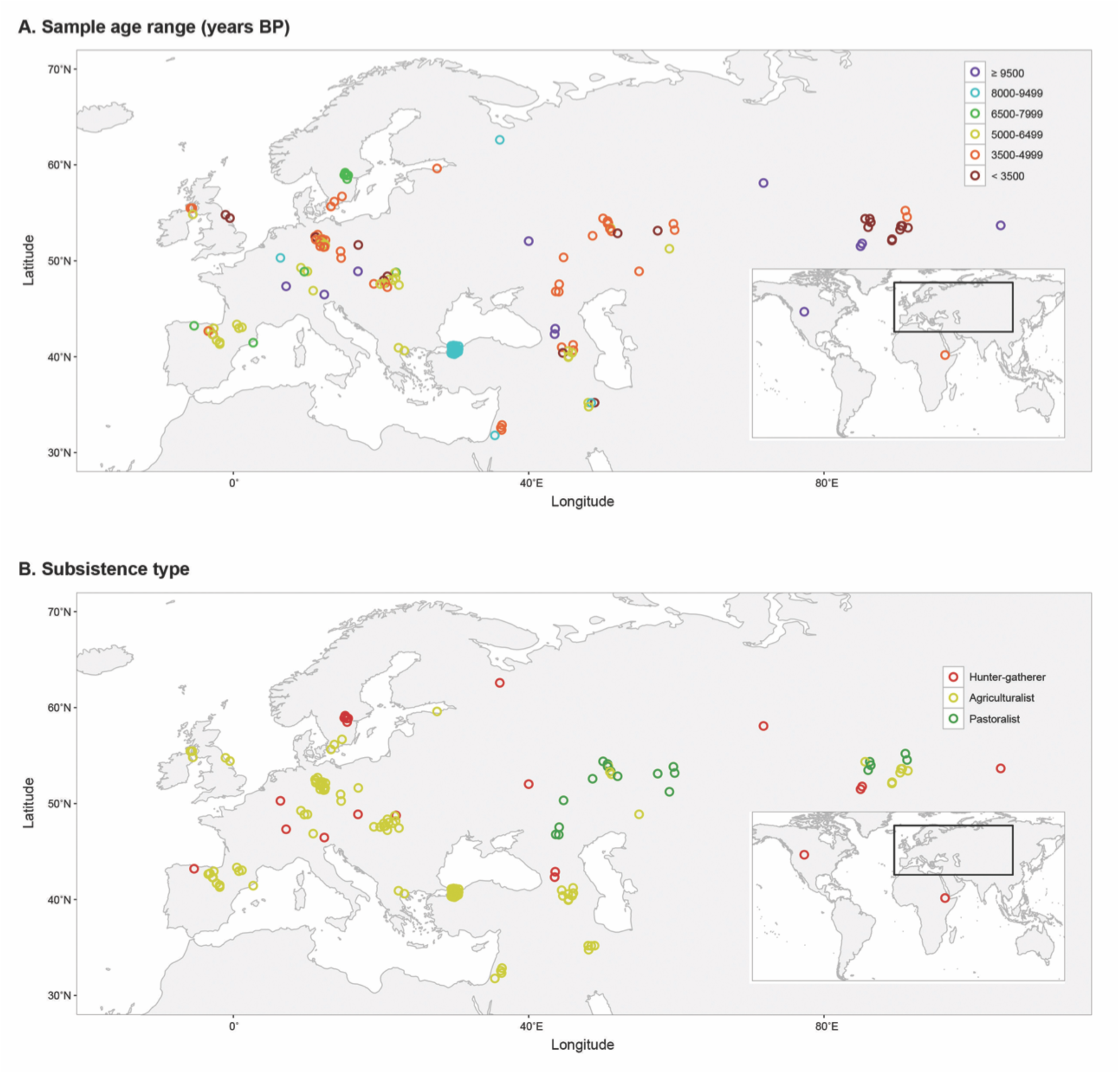
Location, age, and subsistence type of ancient hominins. Geographic distribution of 147 ancient individuals by (A) sample age range in years BP and (B) subsistence type. Over 50% of all disease-associated variants were successfully called for each individual, and geographical locations were jittered to clarify densely sampled locations.

### Modern human genomes

We downloaded modern human genomes from phase 3 of the 1000 Genomes Project (The 1000 Genomes Project Consortium 2015), and we integrated this genomic data with our 3180 focal disease-associated loci. At each of the 3180 loci, we obtained the allele frequencies for the five continental super-populations of modern humans from the 1000 Genomes Project (AFR, AMR, EAS, EUR, and SAS).

### Genetic risk scores

For each ancient sample, we estimated GRS across all diseases and for each disease category by combining genotypic information and effect sizes. All ancient hominins were missing genotypic information at some of the disease-associated loci. To account for this missing information, we calculated GRS for each ancient individual (*i*) from the set of disease-associated loci with genotypic calls in that individual (*L*_*i*_):

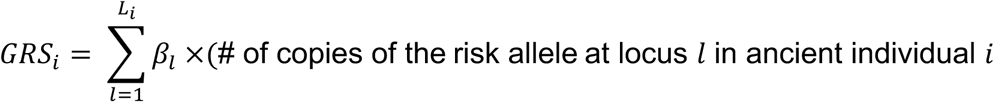

where *β*_*l*_ = *ln* (odds ratio at locus *l*). Because each ancient sample contains a different set of disease-associated loci with genotype calls, GRS are not directly comparable across individuals.

To enable comparisons across individuals, we converted raw GRS statistics into standardized GRS percentiles (see Figure 2 for a pictorial representation). For each ancient genome, this involved simulating 100,000 modern individuals at a matched set of disease loci. Genotypes at disease loci were simulated for 20,000 modern individuals for each of the five super-population from the 1000 Genomes Project (AFR, AMR, EAS, EUR, and SAS). Each diploid genotype was drawn from a binomial distribution using the ‘rbinom’ function in R where the probability of drawing each risk allele is equal to the allele frequency for the given super-population. These simulations assume that disease loci are independent (i.e. they are in linkage equilibrium). Sites that were uncalled in the focal ancient genome were masked in the simulations of modern genomes. Genetic risk scores were then calculated for each modern individual (*J*) using a matched set of called loci (*L*_*i*_) for each focal ancient individual (*i*):

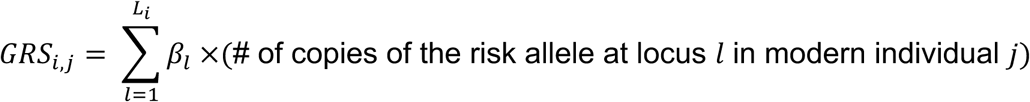

Then, we compared the genetic risk score of each ancient individual to the distribution of the simulated modern genetic risk scores using the empirical cumulative density function (ecdf) function in R. This enabled us to obtain the percentile rank of each ancient sample relative to modern individuals, i.e. the standardized GRS percentile of each ancient genome. We repeated this standardization process for each ancient genome, then used the standardized GRS percentiles to compare genetic risk amongst ancient individuals and between ancient and modern hominins. Higher percentiles indicate less healthy genomes. Standardized GRS percentiles were calculated for the set of all combined diseases and for each disease category. Ancient genomes with risk scores below the range of their matched set of modern genomes have standardized GRS percentiles of 0%, and ancient genomes with risk scores above the range of their matched set of modern genomes have standardized GRS percentiles of 100%. An R Shiny (Chang, et al. 2017) interactive web application of our results can be found at: http://popgen.gatech.edu/ancient-health/.

**Figure 2:**
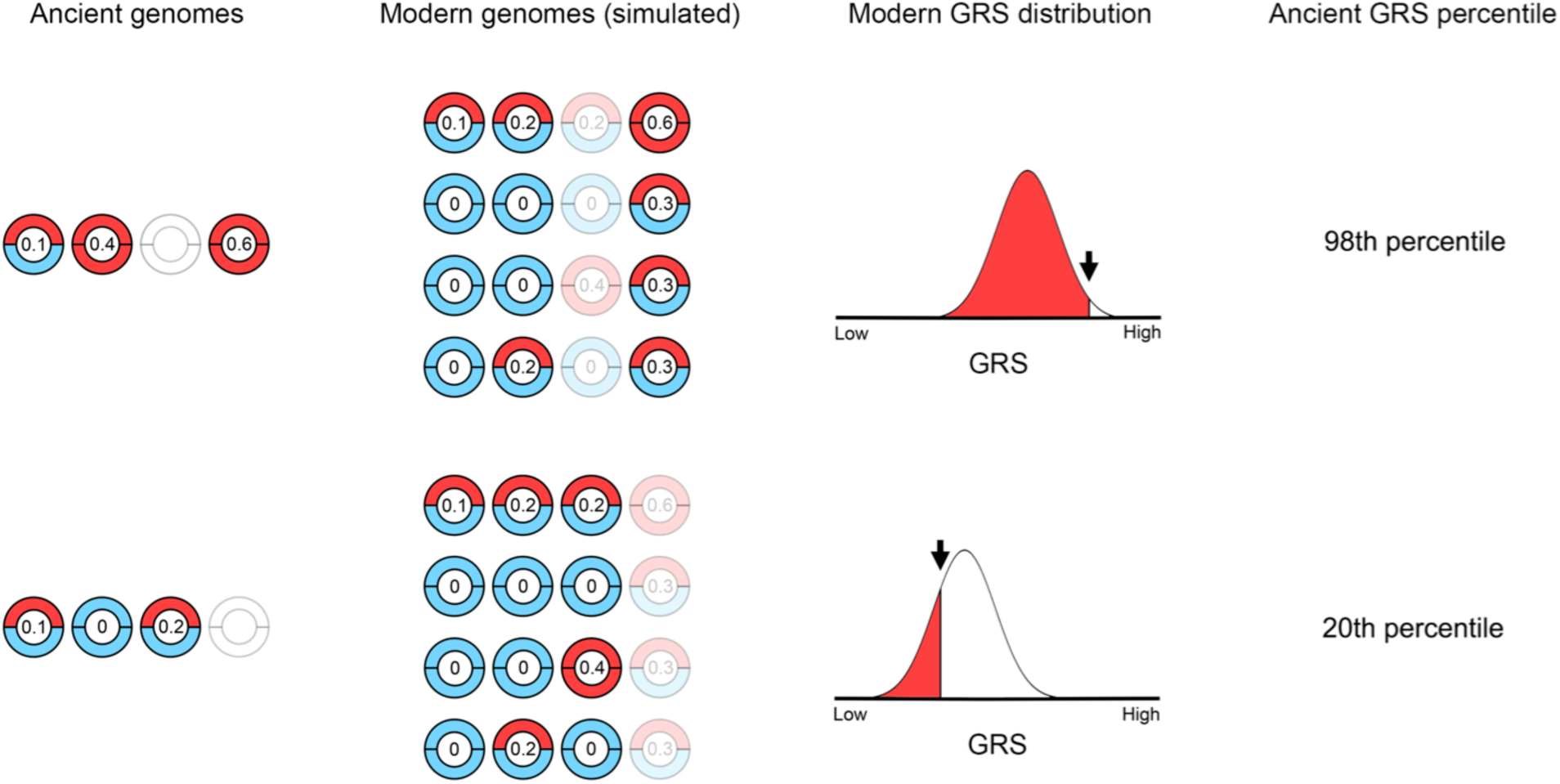
Schematic of how standardized GRS percentiles were estimated for each ancient sample. PokéPlots are shown for each disease locus. Risk alleles are indicated by a red hemisphere, protective alleles are indicated by a blue hemisphere, and numbers denote beta coefficients. Disease loci without genotypic information in the focal ancient individual are masked when estimating GRS for simulated modern humans. Ancient GRS are compared to the distribution of the simulated modern GRS to determine the ancient standardized GRS percentile. The simulated modern GRS distribution differs for each ancient hominin based on which disease loci have known genotypic information.

### Statistical tests

For the set of all disease-associated SNPs and each disease category, we tested for differences between the GRS of modern humans and ancient hominins. By definition, the GRS percentiles of modern humans follow a uniform distribution that ranges between zero and one. If ancient hominins have the same genomic risk of disease as modern humans, the standardized GRS percentile of ancient samples will also be uniformly distributed. We used a Kolmogorov-Smirnov test to determine whether the distribution of ancient hominin standardized GRS percentiles is, in fact, uniformly distributed. With Wilcoxon signed rank tests and continuity corrections, we also tested for a shift in the ancient hominin standardized GRS percentile distribution compared to the expectation of the modern human distribution (centered around 0.5).

Amongst the ancient hominin samples, we investigated whether there are relationships between standardized GRS percentile and sample age, mode of subsistence, and geographic location. These tests were run for all disease loci (overall GRS percentiles) and by disease category. We separated ancient samples into six sample age bins (n = number of samples; < 3500 [n = 20], 3500–4999 [n = 48], 5000–6499 [n = 34], 6500–7999 [n = 10], 8000–9499 [n = 24], and ≥ 9500 years before present [n = 11]). We then tested whether there is a difference in standardized GRS percentile based on sample age bin using a Kruskal-Wallis test. The Kruskal-Wallis test is a rank-based nonparametric test that can be used to determine if the differences between two or more groups are significant. Following rejection of this test, we performed post-hoc multiple pairwise comparisons using Dunn tests to determine which ages were significantly different from each other. Dunn tests generate z-statistics from rank sums. The Benjamini-Hochberg method was used to correct for multiple comparisons. We also used a Jonckheere-Terpstra test, similar to Kruskal-Wallis except that the alternative hypothesis is ordered, to determine whether standardized GRS percentile increased with sample age bin. Mode of subsistence varies across ancient samples (n = number of samples; agriculturalist [n = 106], hunter-gatherer [n = 22], and pastoralist [n = 19]). Differences in standardized GRS percentiles due to mode of subsistence were assessed by Kruskal Wallis tests followed by post-hoc pairwise multiple comparisons with Dunn tests. These non-parametric tests do not require groups to have equal sample sizes. Using Pearson’s correlation, we also tested for an association between standardized GRS percentile and North-South (latitude) or East-West (longitude) gradients.

## Results

### Overall genetic risks

On the whole, ancient samples had GRS that were similar to individuals living in the present (Figure 3A; Supplemental Table S1). Figure 3A shows that the distribution of standardized GRS percentiles of ancient individuals spans the full range of modern humans. In general, most GRS scores are evenly distributed. However, there is an excess of ancient hominins who have standardized GRS percentiles below 5%, which causes the distribution of standardized GRS percentiles to be positively skewed (Kolmogorov-Smirnov test, D = 0.265, p-value = 2.04x10^−9^). Using a Wilcoxon signed rank test, we find that the ancient samples have, on average, lower predicted genomic risk than modern humans (Table 1). However, GRS calculations for ancient samples may underestimate the total amount of genetic risk, as many disease alleles that segregated in the past remain undiscovered.

**Figure 3:**
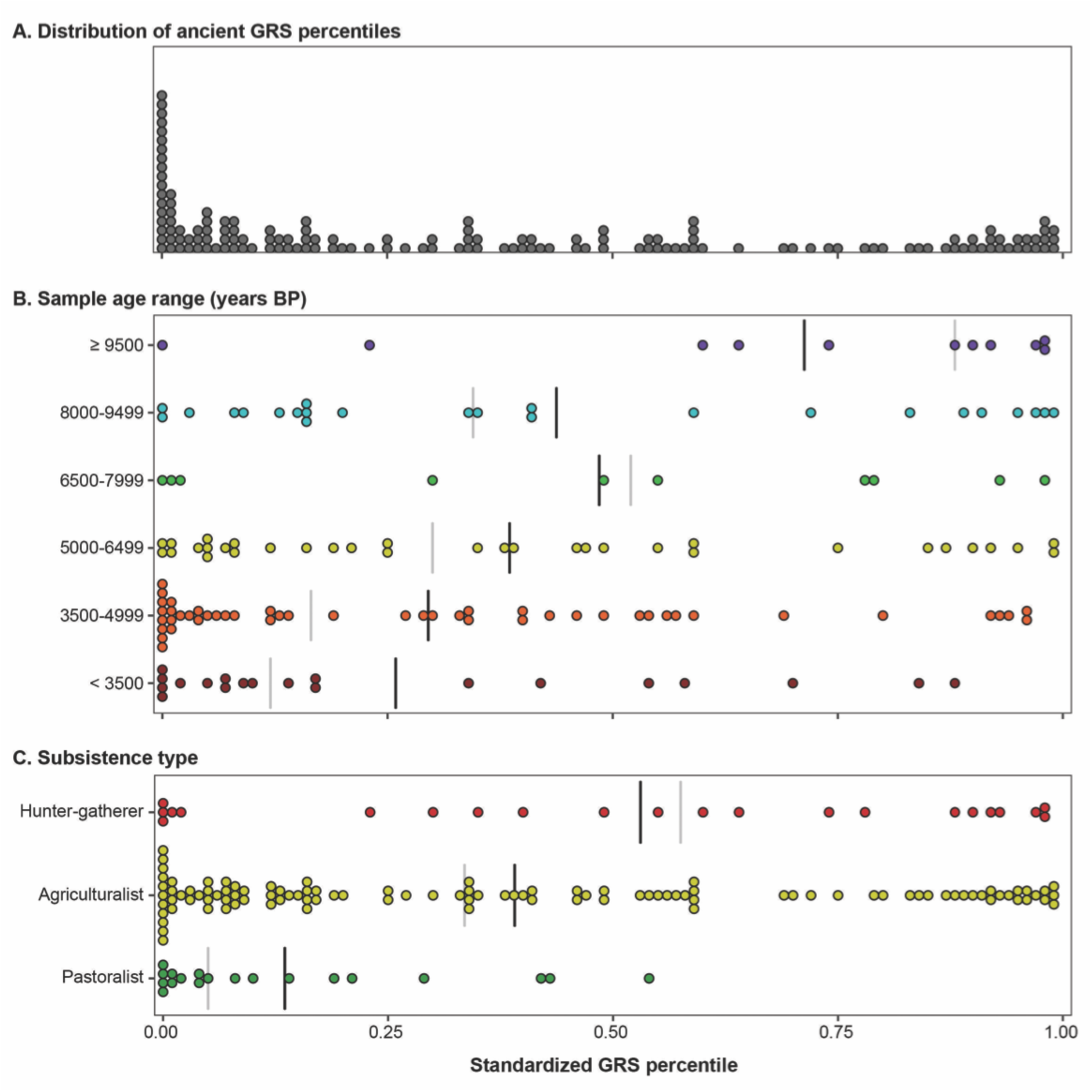
Ancient genetic risk scores span the full range of modern humans. (A) The distribution of ancient standardized GRS percentiles, which includes an excess of healthy individuals below the 5^th^ percentile. Ancient standardized GRS percentiles separated by (B) age range and (C) subsistence type. Group means indicated by black line and group medians denoted by grey line.

**Table 1.** Results of statistical tests between ancient hominin genetic risk scores compared to modern humans (Wilcoxon signed rank test) and amongst ancient samples by sample age range (years BP; Jonckheere-Terpstra test), mode of subsistence (Krustal-Wallis test), and geographic location (latitude and longitude; Pearson’s correlation tests). Boldface indicates that the p-value is less than 0.05.

| Disease Type<br>(number of loci) | Relative to present-day |  | Sample age range |  | Subsistence |  | Latitude |  |  | Longitude |  |  |
| --- | --- | --- | --- | --- | --- | --- | --- | --- | --- | --- | --- | --- |
| | V | p-value | JT | p-value | $\chi^2$ | p-value | r | t | p-value | r | t | p-value |
| All Diseases (3180) | 3266 | **$2.6 \times 10^{-5}$** | 5275 | **$1.0 \times 10^{-3}$** | 12.34 | **$2.0 \times 10^{-3}$** | -0.23 | -2.86 | **$5.0 \times 10^{-3}$** | -0.13 | -1.61 | 0.11 |
| Allergy/Autoimmune (770) | 5028 | 0.51 | 4775.5 | **0.035** | 7.41 | **0.02** | -0.06 | -0.69 | 0.49 | -0.09 | -1.09 | 0.28 |
| Cancers (644) | 4083 | **0.03** | 5049 | **$2.0 \times 10^{-3}$** | 9.73 | **0.01** | -0.29 | -3.71 | **$3.0 \times 10^{-4}$** | -0.08 | -0.94 | 0.35 |
| Cardiovascular (232) | 7659.5 | **$1.8 \times 10^{-5}$** | 4523.5 | 0.18 | 3.26 | 0.20 | -0.06 | -0.70 | 0.49 | -0.20 | -2.44 | **0.02** |
| Dental/Periodontal (40) | 6203 | 0.10 | 4310.5 | 0.38 | 14.15 | **$8.5 \times 10^{-4}$** | -0.20 | -2.48 | **0.01** | -0.13 | -1.57 | 0.12 |
| Gastrointestinal/Liver (488) | 4553 | 0.09 | 4770 | **0.033** | 17.82 | **$1.4 \times 10^{-4}$** | -0.15 | -1.83 | 0.07 | -0.12 | -1.49 | 0.14 |
| Metabolism/Weight (309) | 5052.5 | 0.54 | 4674 | 0.054 | 2.29 | 0.32 | -0.13 | -1.58 | 0.12 | 0.03 | 0.42 | 0.68 |
| Miscellaneous (532) | 3228 | **$1.9 \times 10^{-5}$** | 4451 | 0.24 | 3.63 | 0.16 | -0.06 | -0.74 | 0.46 | $-3.2 \times 10^{-3}$ | -0.04 | 0.97 |
| Morphological/Muscular (239) | 5898 | 0.38 | 4291.5 | 0.42 | 3.62 | 0.16 | 0.14 | 1.70 | 0.09 | -0.14 | -1.66 | 0.10 |
| Neurological/Psychological (459) | 2823 | **$1.8 \times 10^{-6}$** | 4451.5 | 0.25 | 2.83 | 0.24 | 0.01 | 0.12 | 0.90 | $2.8 \times 10^{-3}$ | 0.03 | 0.97 |

Data quality of the ancient genomes varies across samples, so we also tested whether there were any systematic biases in GRS calculations due to sample quality. Both sequencing coverage and the proportion of disease loci that are called may potentially affect whether an individual has a high or low GRS. Based on Pearson’s correlation, there is a weak, yet significant, positive relationship between standardized GRS percentiles and sequencing coverage for ancient samples (Figure 4A, R^2^ = 0.087, p-value = 3.0x10^−4^). However, this pattern may be driven by a small number of outliers, including the Altai Neandertal genome (sequencing coverage: 52x, standardized GRS percentile: 97%). Encouragingly, standardized GRS percentiles are largely independent of the proportion of disease-associated loci that are able to be successfully called in each ancient sample (Figure 4B, R^2^ = 0.013, p-value = 0.17). Taken together, these patterns indicate that our GRS calculations are largely free of biases due to sample quality.

**Figure 4:**
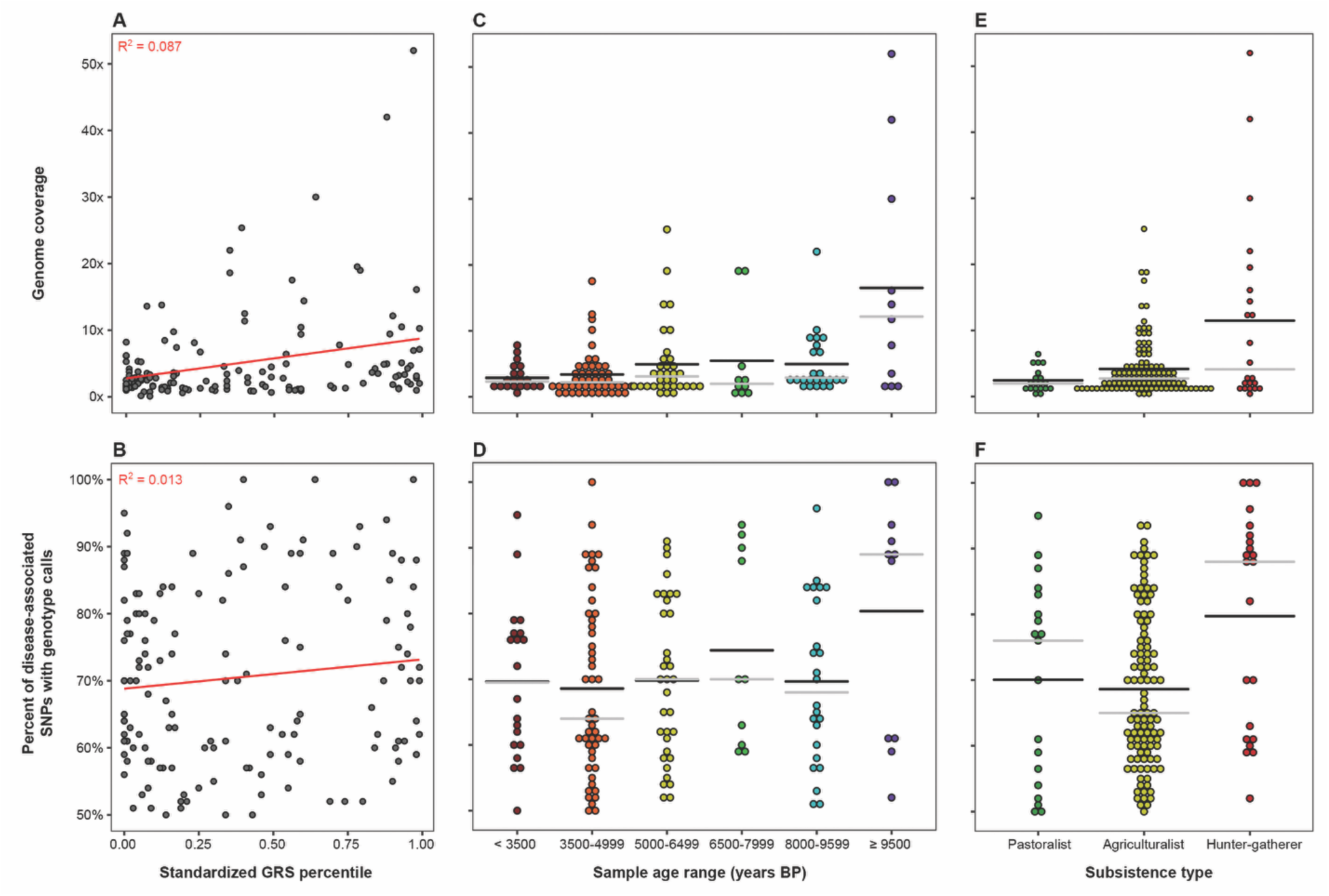
Sequence coverage and sample quality have minimal effect on genetic risk scores. Genome coverage (top) and proportion of disease-associated loci called (bottom) by genetic risk percentile (left), age range (center), and subsistence type (right). Group means and medians indicated by black and grey lines, respectively.

#### Effects of sample age

Figure 3A clearly shows that disease risk varies greatly amongst ancient hominins, so we investigated whether genetic risk scores follow a temporal pattern. Samples were binned into six age ranges, the most recent of which includes individuals that lived less than 3500 years before present and the oldest of which includes individuals who lived at least 9500 years before present (BP). Among ancient samples, there is a difference in the standardized GRS percentile based on sample ages (Kruskal-Wallis test, *χ^2^ =* 15.24, df = 5, p-value = 0.009). From Figure 3B, it is evident that the oldest samples are at much greater genetic risk than more recent samples. We compared standardized GRS percentiles between pairs of sample age ranges using Dunn tests with Benjamini-Hochberg corrections and found that the oldest age range (≥ 9500 years BP) is significantly higher than the two most recent age ranges (< 3500 and 3500–4999 years BP, p-values < 0.007). Following up on this result, we examined whether genetic risks progressively improved with time. Results of a Jonckheere-Terpstra test appear to support this proposition (p-value = 0.001; Table 1), as genetic disease risk increased with later age ranges. This temporal trend is not due to differences in sequence coverage. Our results are robust to the inclusion or exclusion of archaic hominins: when Neanderthal and Denisovan samples are removed, the oldest age class has a median GRS percentile of 0.88.

#### Subsistence type

The number of risk alleles in each ancient genome may also vary by the mode of subsistence, i.e. whether individuals practiced a hunter-gatherer, agricultural, or pastoral lifestyle. We tested this hypothesis by performing a Kruskal-Wallis test. Comparisons of ancient standardized GRS indicate that form of subsistence affects estimated genetic risk of disease (Table 1). Following up on this result, we used a post-hoc Dunn test with Benjamini-Hochberg correction to determine which types of subsistence are different from each one another. Pastoralists have significantly lower standardized GRS percentiles than agriculturalists and hunter-gatherers, while the standardized GRS percentiles are not significantly different between agriculturalists and hunter-gatherers (Table 2). Figure 3C shows that the distribution of standardized GRS percentiles for ancient pastoralists is shifted into to the healthy range of modern humans. We find that the genomes of ancient pastoralists have standardized GRS percentiles almost exclusively below the 50^th^ percentile of modern humans. In contrast, predictions of disease risk in the genomes of ancient agriculturalists and hunter-gatherers span the full range of modern humans.

**Table 2.** Statistical results of ancient genetic risk score comparisons between subsistence types (AG – Agriculturalist; PA – Pastoralist; HG – Hunter-Gatherer) from Dunn tests with Benjamini-Hochberg correction, which were performed on disease categories with significant differences between subsistence groups (see Table 1). Boldface indicates that the p-value is less than 0.05.

|  | AG vs HG |  | AG vs PA |  | HG vs PA |  |
| --- | --- | --- | --- | --- | --- | --- |
|  | z | p-value | z | p-value | z | p-value |
| All Diseases | -1.35 | 0.09 | 2.98 | **$2.1 \times 10^{-3}$** | 3.38 | **$1.1 \times 10^{-3}$** |
| Allergy/Autoimmune | -1.40 | 0.08 | 2.08 | **0.03** | 2.70 | **0.01** |
| Cancers | 0.90 | 0.18 | 3.09 | **$3.0 \times 10^{-3}$** | 1.78 | 0.06 |
| Dental/Periodontal | 3.04 | **$3.5 \times 10^{-3}$** | 2.67 | **$5.7 \times 10^{-3}$** | -0.15 | 0.44 |
| Gastrointestinal/Liver | 0.98 | 0.16 | 4.21 | **$<1.0 \times 10^{-3}$** | 2.61 | **$6.8 \times 10^{-3}$** |

One possible concern is that sequencing coverage and the proportion of disease-associated loci that were successfully called vary by subsistence type (Kruskal-Wallis tests, p-values = 0.03 and 0.01, Figure 4E and 4F, respectively). Differences in sequencing coverage between pastoralists and hunter-gatherers appear to be driven by three older hominin samples – Altai Neandertal (52x), Ust’-Ishim (42x), and Denisovan (30x). However, these differences in sequencing coverage do not lead to significant differences in the proportion of disease-associated loci that are called in the genomes of pastoralists and hunter-gatherers.

#### Geographic patterns

We tested whether genetic estimates of disease risks vary geographically (Table 1). Comparing standardized GRS percentiles and latitude, we find that northern ancient individuals tend to have healthier genomes (Pearson correlation test, *r* = −0.23, p-value = 5.0 × 10^−3^). Interpretation of these results should be tempered by two considerations: 1) most of the ancient samples with sequence data were discovered in Eurasia and 2) there is substantial heterogeneity in genetic risk scores across space. In contrast to latitudinal patterns, the relationship between standardized GRS percentile and longitude does not reach statistical significance (*r* = −0.13, p-value = 0.11). Despite heterogeneity in standardized GRS percentiles and limited sampling, our results suggest that genetic disease risks may have followed a North-South geographic cline in the past.

### Disease Categories

Which types of diseases drive differences in genetic risk scores between ancient and modern individuals? To address this question, we also estimated genetic risk of ancient hominins for nine disease categories (Supplemental Table S1). On average, ancient individuals had significantly lower genetic risk of cancers, miscellaneous diseases, and neurological/psychological diseases than modern humans (Figure 5A; p-values < 0.05). Table 1 lists summary statistics and p-values from Wilcoxon signed ranked tests for each disease category. Only cardiovascular-associated diseases were predicted to have greater risk in ancient hominins compared to modern humans (p-value = 1.8x10^−5^). Risks of allergy/autoimmune, morphological/muscular, metabolism/weight, and dental/periodontal diseases were not significantly different between ancient and modern hominins (p-values > 0.05). Thus, the genetic disease burden of ancient hominins appears to be less than modern humans due to reduced risk of cancer and neurological disease along with other unclassified diseases.

**Figure 5:**
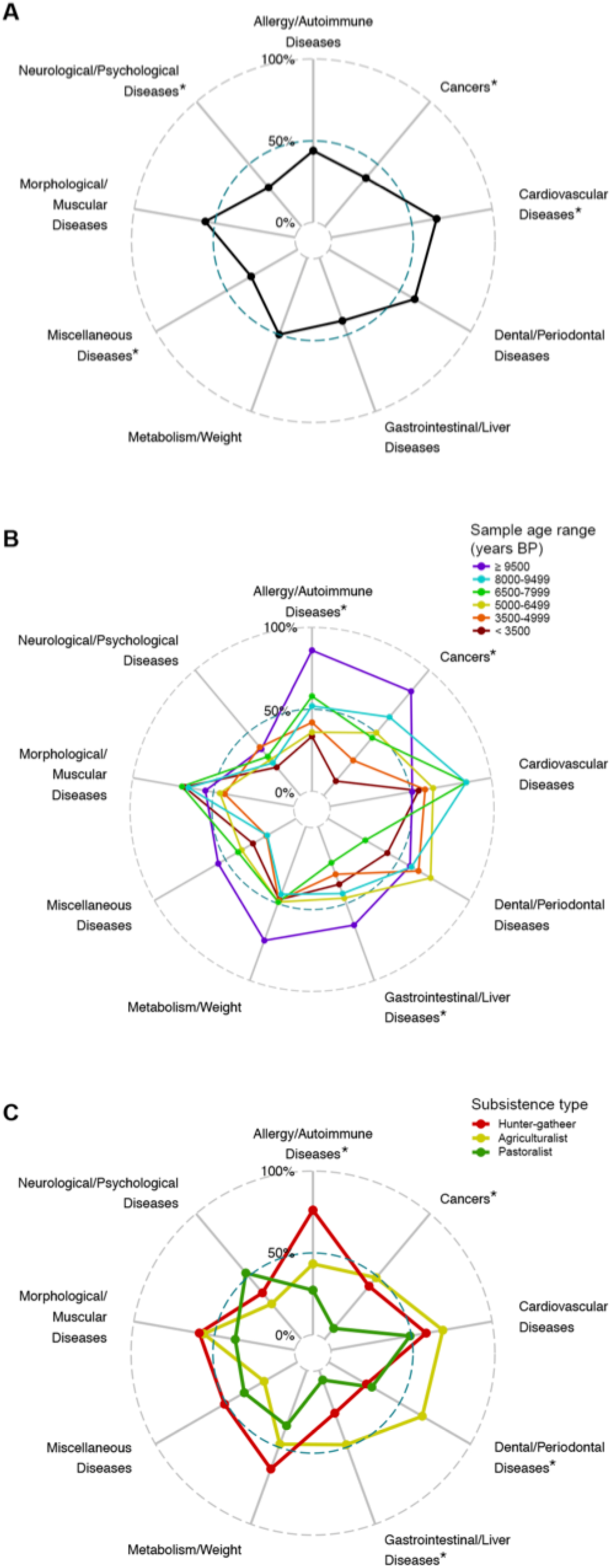
Risk radars of genomic health for ancient samples. The length of each spoke corresponds to a standardized GRS percentile. Healthy individuals have smaller shapes in risk radar plots. (A) Standardized GRS percentile by disease category for all ancient hominins summarized as median risk score. The median of genetic disease risk in modern humans is the 50^th^ percentile. Temporal (B) and subsistence (C) trends of median ancient genetic risk by disease category. Disease categories with significant differences (p-value < 0.05) from Table 1 are denoted by *.

#### Effects of sample age with respect to disease categories

We calculated genetic risks for nine disease categories to determine whether temporal trends were consistent across each disease type. Figure 5B displays the standardized GRS percentile by disease category summarized for each sample age range. Broadly speaking, we find that genetic disease risks change over time. Some disease categories show progressive changes over time. Using Jonckheere-Terpstra tests, we find that older samples have higher standardized GRS percentiles for allergy/autoimmune diseases, cancers, and gastrointestinal/liver diseases (p-values < 0.05). Temporal trends for metabolic diseases were marginally significant (p-value = 0.054). See Table 1 for complete results of Jonckheere-Terpstra tests by disease category. Taken together, the time-dependent results suggest that more recent ancient hominins have lower disease load due to reduced risk of immune-related and gastrointestinal diseases, cancer, and metabolism-related disorders.

#### Subsistence type and disease categories

We tested whether differences in genetic risk for ancient individuals with different modes of subsistence were consistent across all disease categories or whether a single disease category drove disparities in genetic risk between subsistence groups. Using Kruskal-Wallis tests, we find that form of subsistence affects predicted genetic risk of allergy/autoimmune diseases, cancers, dental/periodontal diseases, and gastrointestinal/liver diseases in ancient individuals (Figure 5C). Table 1 includes full results from the Kruskal-Wallis test for each disease category. For each significant disease category, we also performed post-hoc Dunn tests with Benjamini-Hochberg correction; the z-statistics and p-values from these tests are displayed in Table 2. Both the allergy/autoimmune and gastrointestinal/liver disease categories (which share many of the same disease-associated loci) show significantly lower genetic risk in pastoralists than agriculturalists and hunter gatherers. Pastoralists also have significantly reduced risk for cancer compared to agriculturalists. Agriculturalists have a higher genetic risk for dental/periodontal diseases than hunter-gatherers and pastoralists. In general, pastoralists possess extremely healthy genomes, especially for cancers and immune-related, periodontal, and gastrointestinal diseases.

#### Geographic patterns and disease categories

Finally, we examined geographic patterns in genetic risk scores for each disease category using Pearson correlation tests (see Table 1 for results). For most disease types, estimates of genetic risk did not follow a geographic cline. However, decreased risk of disease in northern ancient individuals appears to be driven by two disease categories: cancer (*r* = −0.29) and dental/periodontal diseases (*r* = −0.20). This negative correlation between latitude and standardized GRS percentile is even stronger amongst Eurasian ancient samples (cancer: *r* = −0.33; dental / periodontal diseases: *r* = −0.22). Focusing on longitude, we estimate that the genetic risk of cardiovascular diseases was lower for ancient individuals in the east compared to the west (*r* = −0.20). This trend is slightly weaker when comparing only individuals from Eurasia (*r* = −0.18).

### Ancient precision medicine

Although there are overarching patterns of risk based on type of subsistence and sample age, each ancient individual has their own unique disease profile. These ancient disease risk profiles are summarized in Table S1 and can be viewed via an interactive web application: http://popgen.gatech.edu/ancient-health/. We found that the Altai Neandertal had poor genomic health, with a standardized GRS of 97%. This was due in part to high risk for immune-related diseases, cancers, gastrointestinal and liver diseases, metabolic-related disorders, and morphological and muscular diseases. However, the Altai Neandertal genome did not have an elevated number of risk alleles for all disease classes – we estimated a low risk for cardiovascular disease and close to average risk for periodontal and miscellaneous diseases. Our estimation of genetic risk of cardiovascular disease for the Tyrolean Iceman supports previous evidence that this ancient man had a predisposition for this disease (Keller, et al. 2012). Additionally, the Tyrolean Iceman had high risk for immune-related diseases, gastrointestinal diseases, and metabolic-related disorders. Despite these elevated risks, we also inferred that the genome of the Iceman was relatively healthy for morphological and neurological diseases.

## Discussion

Integrating ancient DNA with knowledge of disease associated loci from modern populations provides new insights into past human health. Overall, we find that the GRS percentiles of ancient hominins span the full range of modern humans. There are some ancient individuals whose genomes place them among the healthiest or least healthy individuals living in the present. Ancient individuals are enriched for standardized GRS percentiles that are below 5% (Figure 3A), which may partially be explained by the fact that our set of known disease-associated loci only includes genetic variants that segregate in modern populations. Thus, we are excluding disease variants that segregated in the past and possibly underestimating the GRS percentile of ancient individuals, i.e. predicted scores are lower than the true genetic disease risk.

The genomic health of ancient individuals appears to have improved over time (Figure 3B). This calls into question the idea that genetic load has been increasing in human populations (Lynch 2016). However, there exists a perplexing pattern: ancient individuals who lived within the last few thousand years have healthier genomes, on average, than present day humans. This deviation from the observed temporal trend of improved genomic health opens up the possibility that deleterious mutations have accumulated in human genomes in the recent past. The data presented here do not provide adequate information to address this hypothesis, which we leave for future follow-up studies.

Genomic studies have shown that there are lingering effects of ancient introgression on the health of modern-day humans. Here, we estimated that the Altai Neandertal genome has an elevated risk of neurological diseases (80^th^ percentile compared to modern humans). This is consistent with previous findings from electronic health records and patient genomes that linked Neandertal haplotypes with increased risk for depression along with tobacco addiction, urinary tract disorders, and skin lesions (Simonti, et al. 2016). Other studies have shown that Neandertal and Denisovan introgression impacted the immune system of present-day humans (Abi-Rached, et al. 2011; Dannemann, et al. 2016; Deschamps, et al. 2016). In our study, the estimated risk of immune-related diseases in Neandertal and Denisovan genomes were at the top of the distribution (above 100% and 97% of all modern genomes, respectively). A previous study showed that archaic alleles at three toll-like receptor genes are associated with reduced microbial resistance and increased risk of allergic diseases (Dannemann, et al. 2016). Despite these ill effects in modern environments, some archaic alleles are quite prevalent in modern Europeans and Asians likely due to natural selection (Deschamps, et al. 2016). For example, there is evidence that significant changes to the coding regions of immunity related genes coincided with the shift in subsistence from hunting and gathering to farming (Deschamps, et al. 2016). Our results suggest that hunter-gatherers had higher genetic risk of allergy and autoimmune disease compared to ancient agriculturalists and pastoralists.

There have been two great dietary shifts in recent human evolution: 1) the consumption of carbohydrate rich diets with the onset of farming (approximately 10,000 years before present) and 2) the utilization of processed flour and sugar since the industrial revolution (approximately 1850 AD). Compared to the other ancient hominins with different forms of subsistence, we estimated that ancient pastoralists had healthier genomes due to low risk for immune-related diseases, gastrointestinal diseases, and cancers. It is unclear why pastoralists would have the lowest risk in these specific disease categories. We caution that this pattern may be the result of technical issues, as pastoralists have the smallest sample size (only 19 individuals) and geographic range (between 40–90°E longitude and 45–55°N latitude, Figure 1B). Because populations that have different subsistence types also differ in other ways, the lower GRS of pastoral populations may be due to other factors, including demographic history.

Comparisons amongst ancient individuals suggest that the transition to agriculture brought about changes not only to diet, but also genetic risk for certain diseases. Based on genomic data, we inferred that ancient agriculturalists had increased risk for dental caries and periodontal diseases. However, slight caution is advised about over interpreting this pattern as dental caries and periodontal diseases had the smallest number of risk loci (k = 40). Previous comparisons of ancient skeletal remains showed that hunter-gatherers had fewer caries than early agriculturalists (Tillier, et al. 1995; Kerr 1998; Trinkaus, et al. 2000; Lebel and Trinkaus 2002). The greater prevalence of dental caries in early agriculturalists may be partially due to differences in oral microbiota diversity, as ancient hunter-gatherers were less likely to have periodontal disease-associated taxa in calcified dental plaques (Adler, et al. 2013).

We note that genomic health does not necessarily equate to phenotypic health. Genetic risk scores are not deterministic, instead they merely indicate whether an individual has a predisposition to a particular disease. In addition, alleles that contribute to disease in modern environments may not have had the same effects in past environments. Because of the large genetic distances between modern humans and archaic hominins, we caution against over-interpreting genomic estimates of Neanderthal and Denisovan health.

Genomic studies, such as this one, provide an opportunity for incomplete or poorly preserved ancient samples to contribute valuable information about the health and disease status of our ancestors. As more ancient hominin genomes are sequenced and new disease-associated loci are identified, there will be unprecedented opportunities to probe changes in human health over time.

